# Sex similarities and dopaminergic differences in interval timing

**DOI:** 10.1101/2023.05.05.539584

**Authors:** Hannah R. Stutt, Matthew A. Weber, Rachael C. Cole, Alexandra S. Bova, Xin Ding, Madison S. McMurrin, Nandakumar S. Narayanan

## Abstract

Rodent behavioral studies have largely focused on male animals, which has limited the generalizability and conclusions of neuroscience research. Working with humans and rodents, we studied sex effects during interval timing that requires participants to estimate an interval of several seconds by making motor responses. Interval timing requires attention to the passage of time and working memory for temporal rules. We found no differences between human females and males in interval timing response times (timing accuracy) or the coefficient of variance of response times (timing precision). Consistent with prior work, we also found no differences between female and male rodents in timing accuracy or precision. In female rodents, there was no difference in interval timing between estrus and diestrus cycle stages. Because dopamine powerfully affects interval timing, we also examined sex differences with drugs targeting dopaminergic receptors. In both female and male rodents, interval timing was delayed after administration of sulpiride (D2-receptor antagonist), quinpirole (D2-receptor agonist), and SCH-23390 (D1-receptor antagonist). By contrast, after administration of SKF-81297 (D1-receptor agonist), interval timing shifted earlier only in male rodents. These data illuminate sex similarities and differences in interval timing. Our results have relevance for rodent models of both cognitive function and brain disease by increasing represenation in behavioral neuroscience.

## INTRODUCTION

The majority of behavioral neuroscience paradigms have been studied using only male rodents, presumably because of variability of the estrus cycle (Beery and Zucker, 2011; Shansky and Woolley, 2016; Shansky, 2019). This has led to biased results that limit the conclusions of neuroscience research. Effects of biological sex and of circulating hormones such as estradiol will help us understand fundamental brain physiology and increase the relevance of basic science research in humans (Bale & Epperson, 2017; Datta et al., 2020; Levy et al., 2023). Notably, human brain diseases can profoundly differ by sex making it an important although understudied biological factor sex (Aleman, Kahn and Selten, 2003; Cahill, 2006; Cerri, Mus and Blandini, 2019).

Understanding sex differences in rodent behavioral tasks is particularly important and highly relevant for rodent models of human brain diseases that display sex differences such as Alzheimer’s disease, Parkinson’s disease, schizophrenia, and substance use disorder, (Abel et al., 2010; Becker & Chartoff, 2019; Gillies et al., 2014; Gür et al., 2019). In humans and rodents, we studied sex differences in interval timing, which has high translational validity for human neurological diseases such as Parkinson’s disease and schizophrenia (C. V. Buhusi & Meck, 2005; Parker et al., 2013; Pastor et al., 1992; Ward et al., 2012). Interval timing involves cognitive processing such as attention to the passage of time and working memory for temporal rules (Rakitin et al., 1998). Our interval timing paradigm requires humans or rodents to estimate an interval of several seconds by making motor responses.

Although the vast majority of prior rodent interval timing studies have used males, a recent report noted no sex differences in interval timing (Buhusi, Bartlett and Buhusi, 2017). In this study we find no differences in interval timing between females and males in humans and rodents, and we found no differences between rodent estrus and diestrus cycle stages. In addition, circulating ovarian hormones such as estradiol can powerfully influence dopamine and dopamine receptors in a sex-dependent manner (Becker, 1990; Meck, 2006). Because interval timing depends on dopamine (Meck, 2006), we explored sex differences in the effect of dopamine receptor pharmacology in interval timing. Our data provide insight into sex similarities and dopaminergic differences during interval timing.

## METHODS

### Human Participants

A total of 45 healthy participants under 50 years old were recruited to perform an interval timing task. Of these participants, 33 were female (median age: 29.1 years; interquartile range (IQR): 16.2 years) and 12 were male (median age: 27.4 years; IQR: 15.5 years). Participants indicated whether they identified as male or female by responding to a questionnaire with the following options: female; male; transgender man/trans man/female-to-male (FTM); transgender woman/trans woman/male-to-female (MTF); genderqueer/gender nonconforming, neither exclusively male nor female; other; and decline to answer. All participants were recruited under IRB #201707828.

### Human interval timing switch task

We adapted an interval timing switch task used previously in rodents and humans (Balci et al., 2009; Larson et al., 2022; Weber et al., 2023). This task required participants to estimate when a short interval (2 seconds) has elapsed by switching response keys before the end of a longer interval (3 seconds; Fig 1A). The trials were displayed on a computer monitor and participants used a standard keyboard to make their responses. There were equal numbers of two types of trials: short interval and long interval. Participants were never told the interval lengths and were trained using the following phases: Phase 1) observe short and long intervals by watching a box appear on the screen for either the short or long interval of time, accompanied by the word “short” or “long” on the screen; Phase 2) observe 10 trials where the box appeared for either the short or long interval and verbally state whether the interval was short or long. They received immediate verbal feedback from the experimenter and had to achieve 80% accuracy to pass this stage; and Phase 3) complete a single 3-minute block of trials during which visual cues indicated accuracy but no points are earned. The testing phase of the task consisted of four 4-minute blocks with ∼60 trials per block.

**Figure 1:**
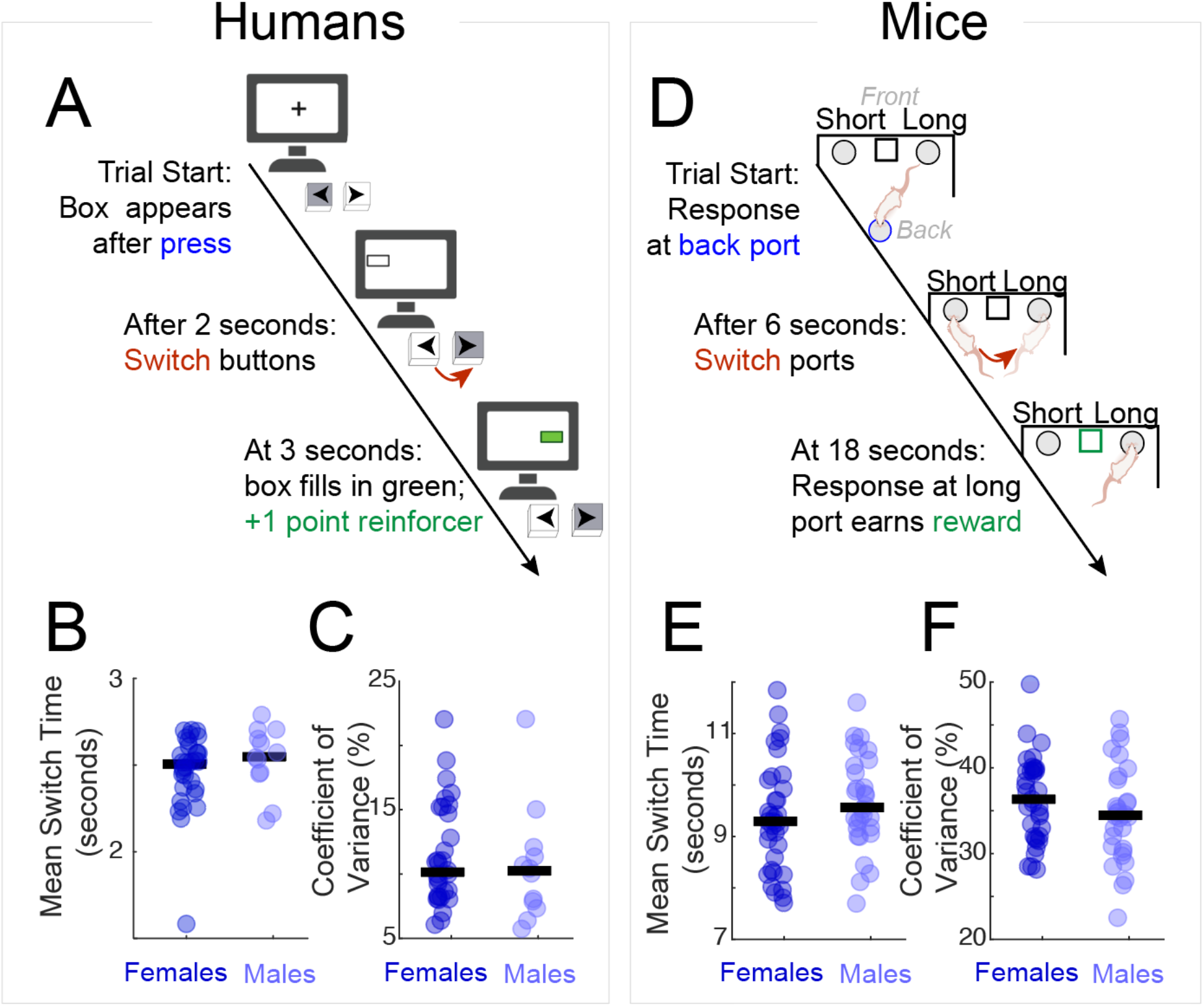
Sex effects on interval timing. A) Human interval timing switch task. The decision to switch buttons is a time-based decision. We measure temporal accuracy by the average switch time and temporal precision by the coefficient of variance of switch times. B) In humans there was no difference in switch times between females and males C) and no difference in the coefficient of variance of switch times between females and males. Data from 33 female and 12 males. D) Rodent interval timing switch task, in which mice switch from a designated “short” response port to a “long” response port after approximately 6 seconds. E) In mice there was no difference in switch times between females and males F) and no difference in the coefficient of variance of switch times between females and males. Solid circles represent the mean switch response or coefficient of variance of switch response time of a single participant. The black horizontal line represents the group median. Data from 35 female and 30 male mice.

All trials started with a white fixation cross displayed on a blank grey screen; the appearance of the white fixation cross cued participants to press and hold the left arrow key, which caused an empty box outlined in green to appear on the left side of the screen. On short-interval trials the box filled in green after 2 seconds and the participant was reinforced with 1 point. If the participant estimated the time interval of 2 seconds had elapsed and the box had not filled in green, the participant then “switched” from pressing the left arrow key to pressing and holding the right arrow key. This switch caused the empty box outlined in green to disappear from the left side and appear on the right side. The empty box on the right filled in green when the long interval of 3 seconds elapsed. If the participant had correctly switched from the left to the right arrow key before the 3 seconds had elapsed, the participant was reinforced with 1 point. If the participant switched before 2 seconds on short interval trials or failed to switch before 3 seconds on long interval trials, the participant earned no points. The decision to switch is guided explicitly by an internal sense of time. Only the data from trials in which a “switch” occurred were analyzed.

### Rodents

All experimental procedures were performed in accordance with Protocol #0062039 and approved by the University of Iowa Institutional Animal Care and Use Committee (IACUC). We included 30 male and 35 female mice. All mice were trained on the same operant-based interval timing task using identical procedures. Although mice had different behavioral endpoints, data were analyzed together because task acquisition and training were identical for all mice. A total of 4 males and 5 females were dopamine-transporter (DAT) IRES-Cre transgenic mice (Jackson #006660; Bar Harbor, ME) maintained on a C57BL/6J background; these mice were bred in-house and had prior stereotaxic surgery. The remaining 26 males and 30 females were C57BL/6J mice ordered from Jackson (#000664). Of these mice, 10 males and 14 females were used to investigate the effects of estrus cycle and dopaminergic drugs. All mice were housed on a 12-hour light-dark cycle with *ad lib* access to water. To facilitate operant behavioral training, all mice were food restricted to ∼85% of free-fed weight. Mice began training on the interval timing task at about 12 weeks of age.

### Rodent interval timing switch task

We trained mice on an interval timing switch task that assessed their ability to control their actions based on their internal representation of time (Fig 1D)(Balci et al., 2008; Larson et al., 2022; Weber et al., 2023). Training occurred in operant chambers placed in sound attenuating cabinets (MedAssociates, St. Albans, VT). Chambers contained two response ports at the front separated by a food hopper, and one response port on the back wall. Nosepoke responses were detected by infrared beams at each port.

A trial was initiated by a nosepoke at the back response ports which triggered cue lights above each front port and an 8-kHz tone at 72 dB. Short- and long-interval trials had identical cues. All trials were reinforced with 20 mg sucrose pellets (BioServ, Flemington, NJ). Short intervals occurred on 50% of trials and were reinforced for the first response after 6 seconds at the designated “short port” (counterbalanced left or right). Long-interval trials were reinforced only when the mouse responded after 18 seconds by “switching” from the short port to the “long port”. On these switch trials, mice responded at the short port until approximately 6 seconds has passedand then switched to the long port until reward delivery for the first response after 18 seconds. During switch trials, when animals depart the last response at the short response port is considered the “switch response time” and measures rodent’s internal estimate of time for when to switch from the short to long response port. Only the data from long-interval switch trials were analyzed. Two experimental sessions of testing data per mouse were collected and analyzed for each treatment condition; sessions with <2 completed switch trials were excluded.

Experimental sessions lasted 90 minutes.

### Drug preparation and administration

We examined how drugs targeting dopaminergic receptors affected interval timing as a function of sex. All drugs were purchased from Tocris Bioscience (Bristol, UK: sulpiride #0894; quinpirole #1061; SCH-23390 #0925; SKF-81297 #1447;). Doses were based on prior literature and pilot experiments (del Arco et al., 2004; Dias et al., 2010; Zarrindast et al., 2008). Sulpiride was dissolved in acidified sterile 0.9% saline (15% acetic acid; pH 2.0), which was then neutralized using NaOH pellets to pH ∼7.1–7.2. SCH-23390, SKF-81297, and quinpirole were dissolved in sterile 0.9% saline.

Once sufficient behavior was achieved during interval timing, mice were tested on drug administration sessions. Before testing sessions, mice were injected intraperitoneally (IP) with either saline, sulpiride (12.5 mg/kg), quinpirole (0.15 mg/kg), SCH-23390 (0.05 mg/kg), or SKF-81297 (males: 0.5 mg/kg; females: 0.5 mg/kg or 1.5 mg/kg), in a pseudorandomized counterbalanced design. The experimenter administering the drugs was blinded to treatment condition.

### Vaginal cytology

In mice, estrus cycle stage was assessed using vaginal cytology. Using a sterile 100-µl pipette tip, 15 µl of phosphate buffered saline (PBS) was inserted in the vagina; the pipette plunger was depressed and retracted twice before applying the sample on a microscope slide. Samples were assessed for estrus cycle stage using a Leica 10x microscope. The estrus stage was identified by the presence of cornified cells and the absence of leukocytes; the diestrus stage was identified by the presence of cells comprised primarily of leukocytes (Byers et al., 2012; Caligioni, 2009).

### Statistics

Interval timing performance was quantified by two metrics: 1) the mean switch time in a session, which is a measure of timing accuracy, and 2) the coefficient of variation (CV) of switch times in a session, which is a measure of timing precision. We also analyzed the number of responses in a session as a metric to ensure task motivation was consistent between groups. The CV is calculated as standard deviation divided by the mean and is expressed as a percentage. All values were compared by nonparametric rank sum or signed rank tests using the *ranksum* or *signrank* functions in MATLAB (R2022b).

## RESULTS

We examined sex differences in interval timing using similar tasks in both humans and rodents (Fig 1A and 1D). In humans, we found similar switch times between females and males (all results presented as median (Q1-Q3); females: 2.5 seconds (2.4–2.6); males: 2.6 seconds (2.5–2.7); p = 0.23). The switch time CVs between females and males were also similar (females: 10% (8%–15%); males: 10% (8%–12%); p = 0.45). In rodents, we also found that female and male mice had similar switch times (females: 9.3 seconds (8.4–10.1); males: 9.6 seconds (9.2–10.4); p = 0.17; Fig 1B) and switch time CVs (females: 36.3% (32.0%–39.7%); males: 34.5% (30.6%–38.6%); p = 0.20; Fig 1C). In addition, female and male mice had a similar number of total responses within each session (females: 440 responses (405–571); males: 495 responses (422–644); p = 0.17; Fig 1E). These data support prior interval timing studies that reported no differences in interval timing with sex in rodents (Buhusi, Bartlett and Buhusi, 2017).

Historical concerns about including female rodents often relate to the assumption that fluctuating ovarian hormones increase behavioral variability (Beery & Zucker, 2011; McCarthy et al., 2012). Because ovarian hormones fluctuate throughout the reproductive cycle, we used vaginal cytology to track the estrus cycle in female mice. We found no significant differences in switch times associated with female mouse reproductive cycles (estrus: 9.3 seconds (8.8–9.9); diestrus: 9.2 seconds (8.7–9.9); p = 0.86; Fig 2A); switch time CVs (estrus: 31.7% (27.7%– 36.7%); diestrus: 33.2% (30.8%–36.5%); p = 0.71; Fig 2B); or number of responses (estrus: 355 responses (287–401); diestrus: 362 responses (324–478); p = 0.09; Fig 2C). These data demonstrate that interval timing is consistent throughout the rodent reproductive cycle.

**Figure 2:**
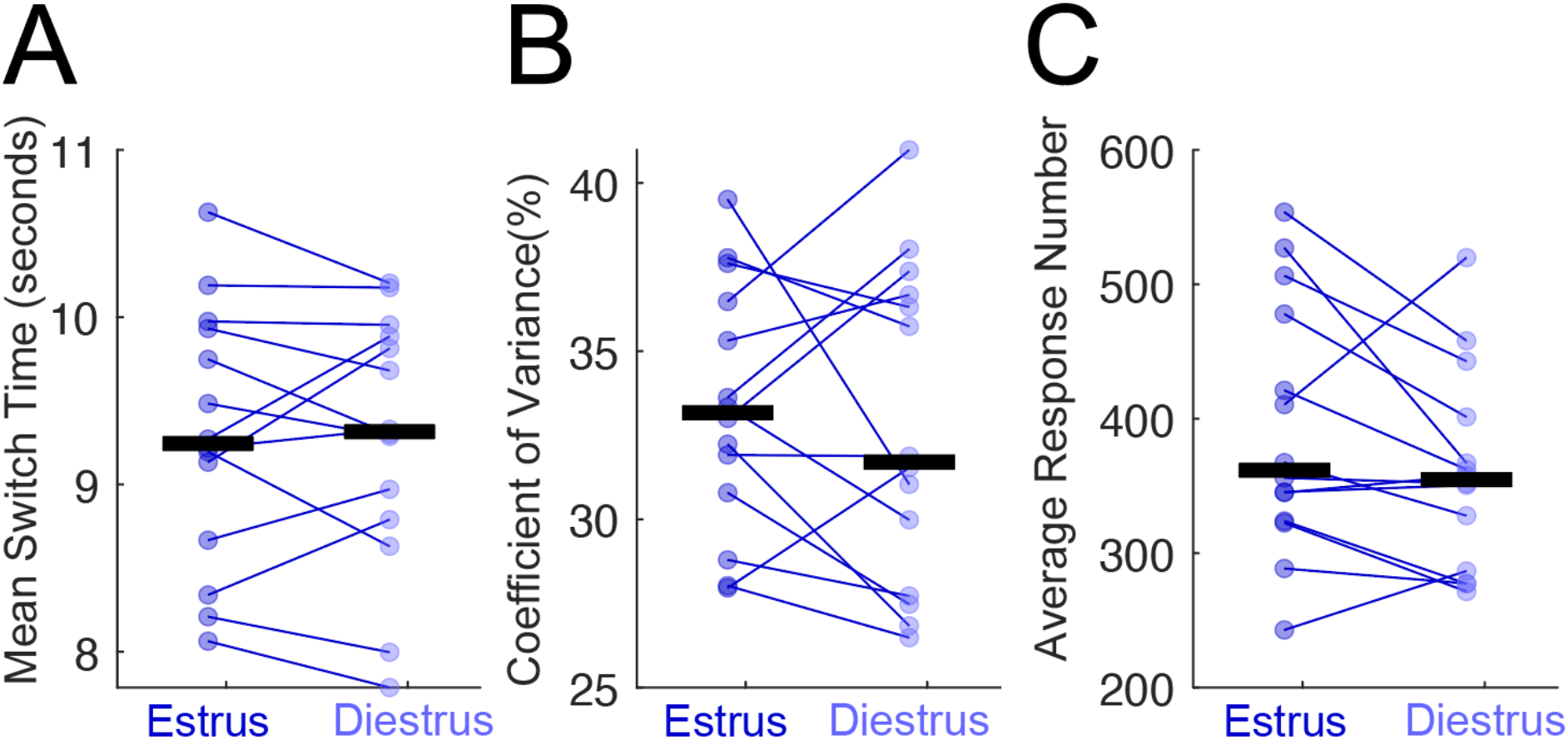
Estrus cycle does not affect interval timing. A) There was no difference in the switch time, B) coefficient of variance, or number of responses between females in the estrus and diestrus stage. Solid circles represent the mean switch response or coefficient of variance of switch response time or average number of responses of a single mouse. The black horizontal line represents the group median. Data from 14 female mice.

Dopamine can powerfully affect interval timing (De Corte et al., 2019; Meck, 2006; Soares et al., 2016; Ward et al., 2009). We examined how drugs targeting dopaminergic receptors affected interval timing as a function of sex. In line with prior work, the D2 antagonist sulpiride delayed switch times in both females (saline: 9.4 seconds (8.6–10.3); sulpiride: 11.1 seconds (9.5–11.5); p = 0.001; Fig 3A) and males (saline control: 8.8 seconds (8.4–9.9); sulpiride: 10.3 seconds (9.7–10.8); p = 0.006). In addition, after sulpiride administration the switch time CV was significantly decreased in both females (saline: 34.2 (31.6–34.8); sulpiride: 28.4 (25.2–29.8); p = 0.003; Fig 3B) and males (saline control: 33.5 (30.9–37.4); sulpiride: 29.8 (27.8–32.9); p = 0.004; Fig 3B). The D2 agonist quinpirole also delayed switch times in females (saline control: 9.2 seconds (8.8–9.5); quinpirole: 10.0 seconds (9.0–10.9); p = 0.0001; Fig 3C) and males (saline control: 8.8 seconds (8.3–9.3); quinpirole: 9.7 seconds (8.9–11.0); p = 0.01; Fig 3C). Quinpirole did not reliably affect CV for females (saline: 30.6 (26.7 - 34.3); quinpirole: 32.1 (25.5 - 33.1); p= 0.43; Fig 3D) or males (saline: 30.4 (27.7 - 34.9); quinpirole: 29.0 (26.8 - 34.3); p=0.13; Fig 3D).

**Figure 3.**
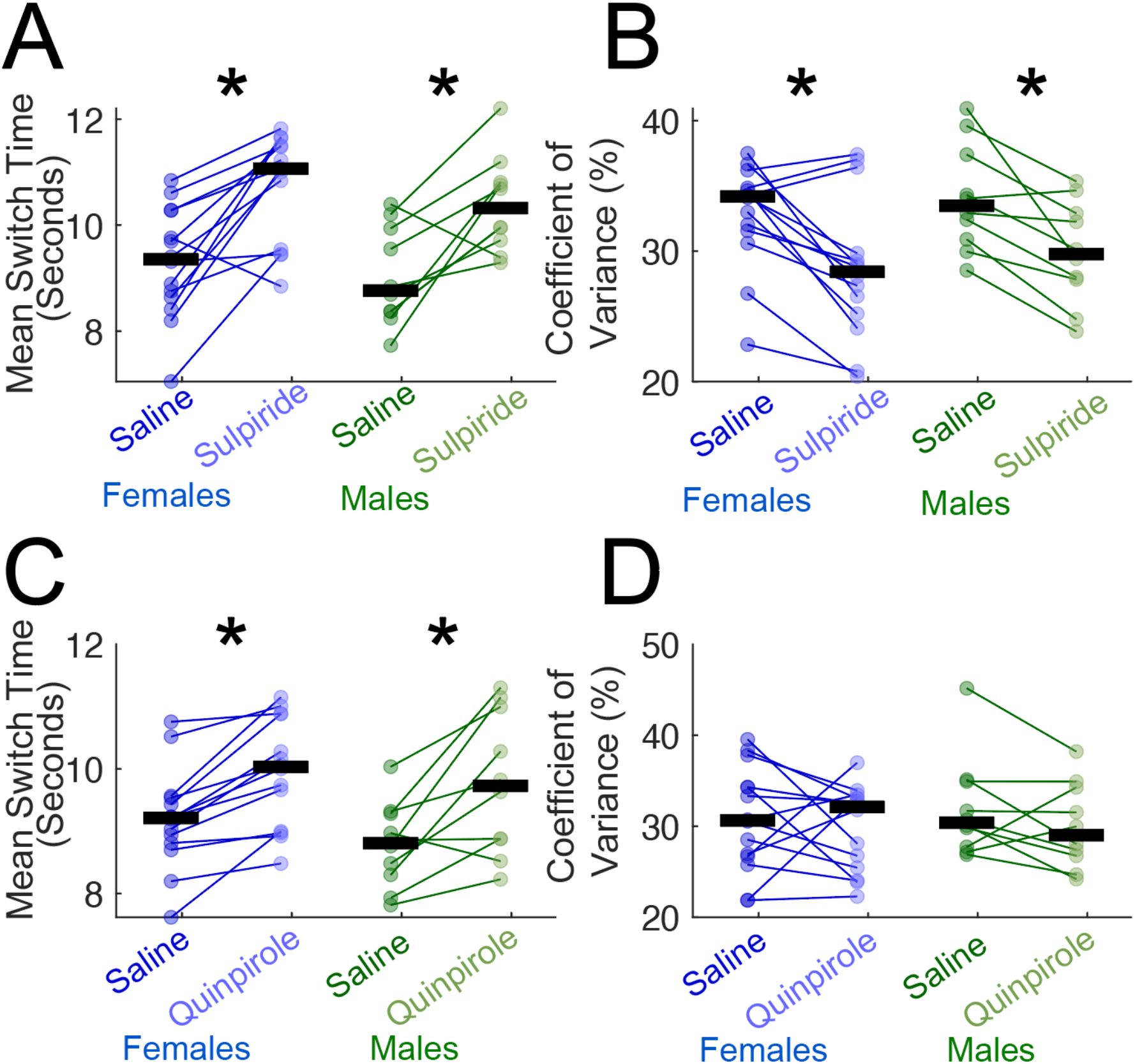
Female and male switch times and coefficient of variance after D2 agonists and antagonists. A) The D2 antagonist sulpiride significantly increased switch times in both females and males. B) Sulpiride also significantly decreased switch time coefficient of variance in both females and males. C) The D2 agonist quinpirole also significantly increased switch times in both females and males but D) did not reliably affect switch time coefficient of variance. Solid circles represent the mean switch response or coefficient of variance of switch response time of a single mouse. The black horizontal line represents the group median. *p < 0.05. Data from 14 females and 10 males.

The D1 antagonist SCH-23390 also delayed switch times in both females (saline control: 9.3 seconds (8.9–9.8); SCH-23390 10.4 seconds (9.6–11.0); p = 0.01; Fig 4A) and males (saline control: 9.1 seconds (8.1–9.3); SCH-23390: 10.4 seconds (9.5–10.7); p = 0.002; Fig 4A). SCH-23390 did not reliably affect CV in either females (saline: 30.9 (27.1 - 31.4); SCH: 28.6 (26.9 - 33.1); p=0.63; Fig 4B) or males (saline: 31.7 (30.6 - 36.5); SCH: 28.5 (26.1 - 30.7); p=0.06; Fig 4B). Interestingly, we found that the D1 agonist SKF-81297 at a dose of 0.5 mg/kg decreased switch times in male mice only (saline control: 8.8 seconds (8.1–9.1); SKF-81297: 9.4 seconds (8.8–9.8); p = 0.02; Fig 4C). Strikingly, there was no difference in switch time for female mice after SKF-81297 was administered at 0.5 mg/kg (saline control: 8.4 seconds (7.8– 9.1); SKF-81297: 9.0 seconds (8.5–9.7); p = 0.30; Fig 4C) or at a higher dose of 1.5 mg/kg (saline control: 8.6 seconds (7.9–9.8); SKF-81297: 8.5 seconds (8.2–9.4); p = 0.67; Fig 4C). In addition, SKF-81297 did not reliably change switch time CVs for males at the 0.5mg/kg dose (saline: 30.0 (28.1 - 31.2); SKF: 31.6 (28.1 - 36.7); p=0.43; Fig 4D) or females at the 0.5mg/kg (saline: 30.4 (29.6 - 39.7); SKF: 33.6 (32.9 - 37.2); p=0.38; Fig 4D) and 1.5mg/kg dose (saline: 33.3 (32.1 - 40.0); SKF: 35.1 (32.7 - 38.4); p=1.00; Fig 4D). The number of responses did not differ at the 0.5 mg/kg dose for males (saline: 487.2 (343.5 - 588.5); SKF: 512.0 (360.0 - 590.5); p=0.62). However, in females the average response number was significantly decreased at the 1.5 mg/kg dose (saline: 378 responses (284–488); SKF-81297: 340 responses (297–380); p = 0.01). but not the 0.5mg/kg dose (saline: 334.5 (329.9 - 433.2); SKF: 341.5 (333.0 - 397.5); p=0.47). Taken together, these data describe sex-dependent similarities and differences in interval timing.

**Figure 4.**
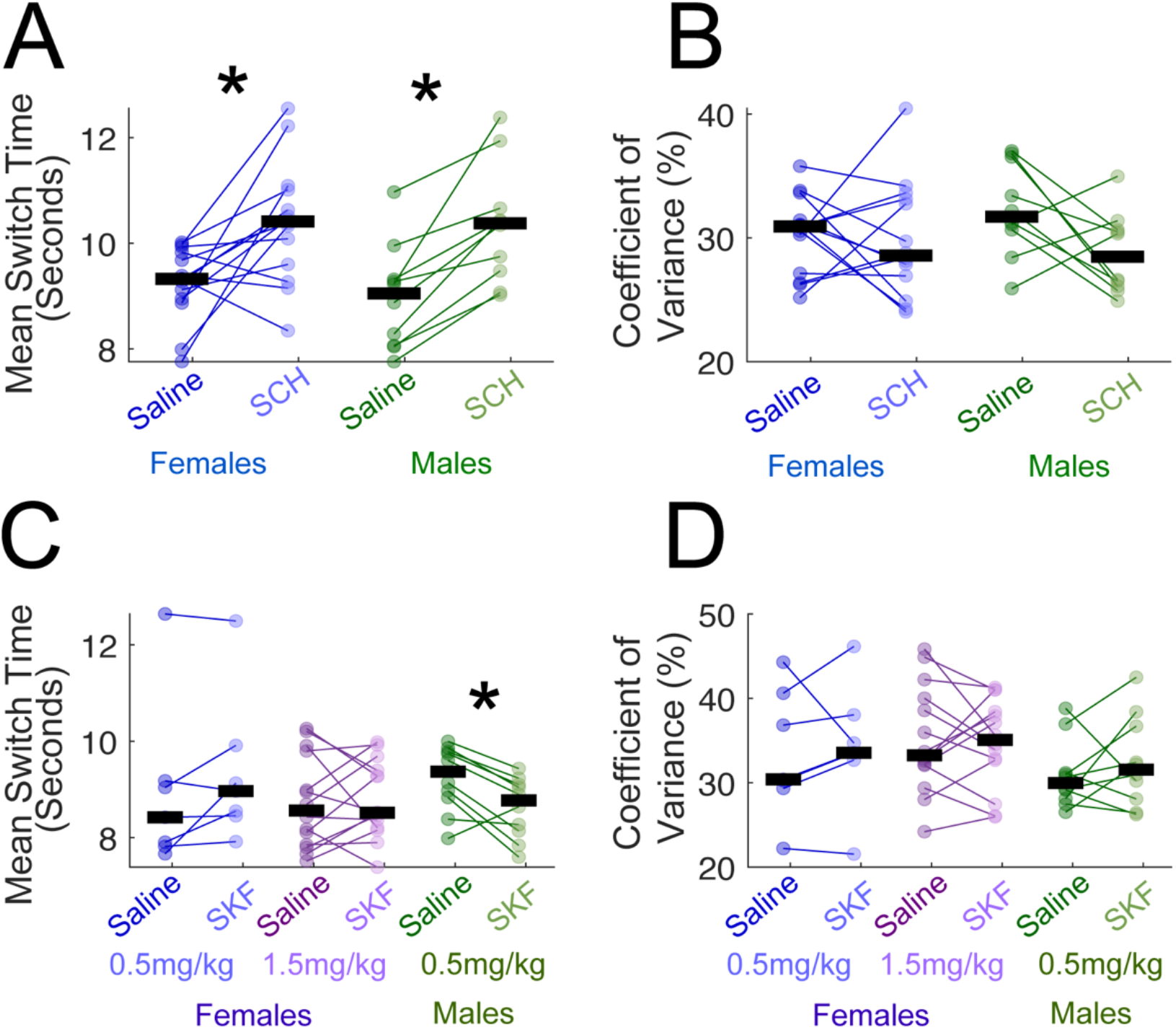
Female and male switch times and coefficient of variance after D1 agonists and antagonists. A) The D1 antagonist SCH-23390 (SCH) increased switch times in both females and males, while B) there was no significant difference in the switch time coefficient of variance in females or males. C) The D1 agonist SKF-81297 (SKF) did not reliably change switch time at doses of 0.5 mg/kg nor 1.5 mg/kg in females. However, SKF at a dose of 0.5 mg/kg dose decreased switch times in males only. D) SKF did not induce any significant difference in switch time coefficient of variance in females or males. Solid circles represent the mean switch response or coefficient of variance of switch time of a single mouse. The black horizontal line represents the group median. *p<0.05. SCH data comes from 14 females and 10 males. SKF at the dose of 1.5mg/kg comes from 14 females, and data from the 0.5mg/kg dose comes from 6 females. 10 males were included in the SKF 0.5mg/kg data.

## DISCUSSION

We demonstrate that temporal accuracy and temporal precision is similar or not different between females and males in humans and mice. We also report that the rodent estrus cycle stage does not influence interval timing. D2 antagonists, D2 agonists, and D1 antagonists have similar effects on interval timing in female and male mice; however, D1 agonists affected timing in male mice only, implicating D1-receptor defined circuits in sex-specific effects. These findings corroborate prior studies that found no sex differences in interval timing performance (Buhusi, Bartlett and Buhusi, 2017). Our results have important implications for addressing claims about the female reproductive cycle introducing experimental variability (Fields, 2014; Levy et al., 2023; Richardson et al., 2015). This study is important for informing future experimental manipulations in cognitive behaviors.

We found sex differences in switch times after administration of the D1 agonist SKF-81297. Male mice had earlier switch times at a dose of 0.5 mg/kg, whereas female mice displayed no difference in switch times at 0.5 mg/kg or the higher dose of 1.5 mg/kg. These findings could indicate a mechanistic sex difference, as prior work in cell culture found that another D1 agonist, SKF-82958, competes with 17*β*-estradiol to bind to estrogen receptor alpha (ER*α*) and beta (ER*β*) (Walters et al., 2002). Both males and females express ER*α* and ER*β*, although there could be sex differences in downstream effects of receptor activation that contribute to this behavioral sex difference. Future studies will elucidate the mechanism of these differences.

Our study has several limitations. First, interval timing measures only some aspects of cognitive function; other assays might be more sensitive to sex differences. Second, mice from the C57BL/6J strain were used, and it is unclear whether our results generalize to other strains.

Further studies using additional cognitive tasks and additional rodent strains will be necessary to determine if females and males differ in cognitive performance.

In conclusion, we found no reliable sex differences in interval timing in humans or rodents, and we determined that the rodent estrus cycle does not affect interval timing. While D2 antagonists, D2 agonists, and D1 antagonists increased switch times for female and male mice, D1 agonists had sex-specific effects and only decreased switch times in male mice. These results help promote the use of both female and male rodents in behavioral neuroscience.

## DATA AND CODE AVAILABILITY

All raw data and code are available for download at narayanan.lab.uiowa.edu/datasets.

## ACKNOWLEDGEMENTS

Funded by NINDS R01 NS120987 and NIMH R01MH116043. We are grateful to Fuat Balci for sharing the human switch task.

## COMPETING INTERESTS

None

## AUTHOR CONTRIBUTIONS

HRS, MAW, ASB, RCC, and NSN designed experiments, HRS, XD, RC and MSM performed experiments; HRS, MAW, and RCC, analyzed data, and HRS, MAW, ASB, RC, and NSN wrote the manuscript.

## Notes

https://narayanan.lab.uiowa.edu/datasets

### Competing Interest Statement

The authors have declared no competing interest.

